# Genetic comparisons of interleukin-17 reveal a framework for complex signaling evolution

**DOI:** 10.64898/2026.04.13.718218

**Authors:** Steve S. Cho, Gloria B. Choi, Jun R. Huh, Nels C. Elde

## Abstract

The interleukin-17 (IL-17) family of cytokines comprises structurally distinct ligands and receptors which mediate immune responses at mucosal surfaces. The growing understanding of its regulatory functions beyond immunity, together with extensive genetic variation in protein-coding genes, raises the possibility that IL-17 cytokines participate in an even wider network of biologic processes. Despite successes of experimental approaches to chart IL-17 functions, inherent signaling complexities and crosstalk with multiple physiologic pathways obscure a full appreciation of the biological potential of IL-17. Here, we integrated comparative genomics, evolutionary rate covariation (ERC), and signatures of natural selection to resolve phylogenetic relationships between IL-17 ligands and receptors and discovered evidence for hidden signaling interactions. ERC analysis revealed putative ligand-receptor interactions for IL-17D and IL-17RC and suggested uncharacterized potential signaling mediator for the receptor IL-17REL, such as IL-17B. Signals of covariation extended beyond the IL-17 family to other genes encoding neurodevelopmental effectors and growth factors, emphasizing recurrent co-evolutionary patterns that delineate the immune and neuromodulatory roles of IL-17. These connections are underlined by signatures of positive selection in the disordered N-terminal domain of IL-17E and its cognate receptor, IL-17RB, key modulators of both type 2 immune response and neuronal function, suggesting functional consequences of this understudied domain. Together, our findings suggest that IL-17 biology is repeatedly impacted by lineage-specific selective pressures that dictate both immune and non-immune functions. By anchoring the expanding IL-17 field in an evolutionary framework, we propose a model for understanding the diversification and functional expansion of this and other cytokine families.

## Introduction

Cytokines are essential orchestrators of the immune response against various stimuli, including pathogens and cancer. While many vertebrate immune effectors can be traced to a single ancestral acquisition event, millions of years of evolution have driven the functional diversification of their orthologs (Margolis, et al. 2017; Daly, et al. 2025; Langley, et al. 2025). Indeed, we observe numerous recurrent adaptations, homolog expansions, and gene losses in cytokines across metazoans (Brocker, et al. 2010). These adaptations in response to both pathogenic and non-pathogenic selective pressures impact biological functions beyond their canonical roles in inflammatory immune regulation (Salvador, et al. 2021; Yang, et al. 2023).

A growing body of evidence suggests that the interleukin-17 (IL-17) cytokine family mediates non-immune functions beyond their inflammatory effector capacities. The IL-17 pathway arose in ancient eumetazoans from a domain-shuffling event between two ancestral signaling pathways, constituting the only cytokine family conserved in both vertebrates and invertebrates (Chen, et al. 2024). This resulted in a cysteine knot receptor-binding domain, a conserved structural motif stereotypically associated with growth factors and neurotrophins, which structurally distinguishes IL-17 ligands from other cytokines (Zhang, et al. 2011). In humans and most mammals, six different IL-17 ligands (IL17A through F) signal as homo- or heterodimers and interact with heterodimeric complexes of six cognate IL-17 receptor (IL-17RA through E, REL) subunits on the cell surface membrane, leading to activation of the NF-κB-mediated inflammatory cascade (**Figure 1A**)(Gaffen 2009; McGeachy, et al. 2019). Despite greater structural homology to neurodevelopmental effectors (e.g. neurotrophins and nerve growth factors) than to other immune-related cytokines, mammalian IL-17A and IL-17F, the two most widely studied IL-17 ligands, are secreted by multiple lymphocyte populations such as T-helper type 17 cells (Th17 cells), γδ T cells, and group 3 lymphoid cells (ILC3)(Sutton, et al. 2012). Accordingly, most functional studies of IL-17 family proteins have been performed in the context of these and similar immune responses (McDonald and Hendrickson 1993).

**Figure 1.**
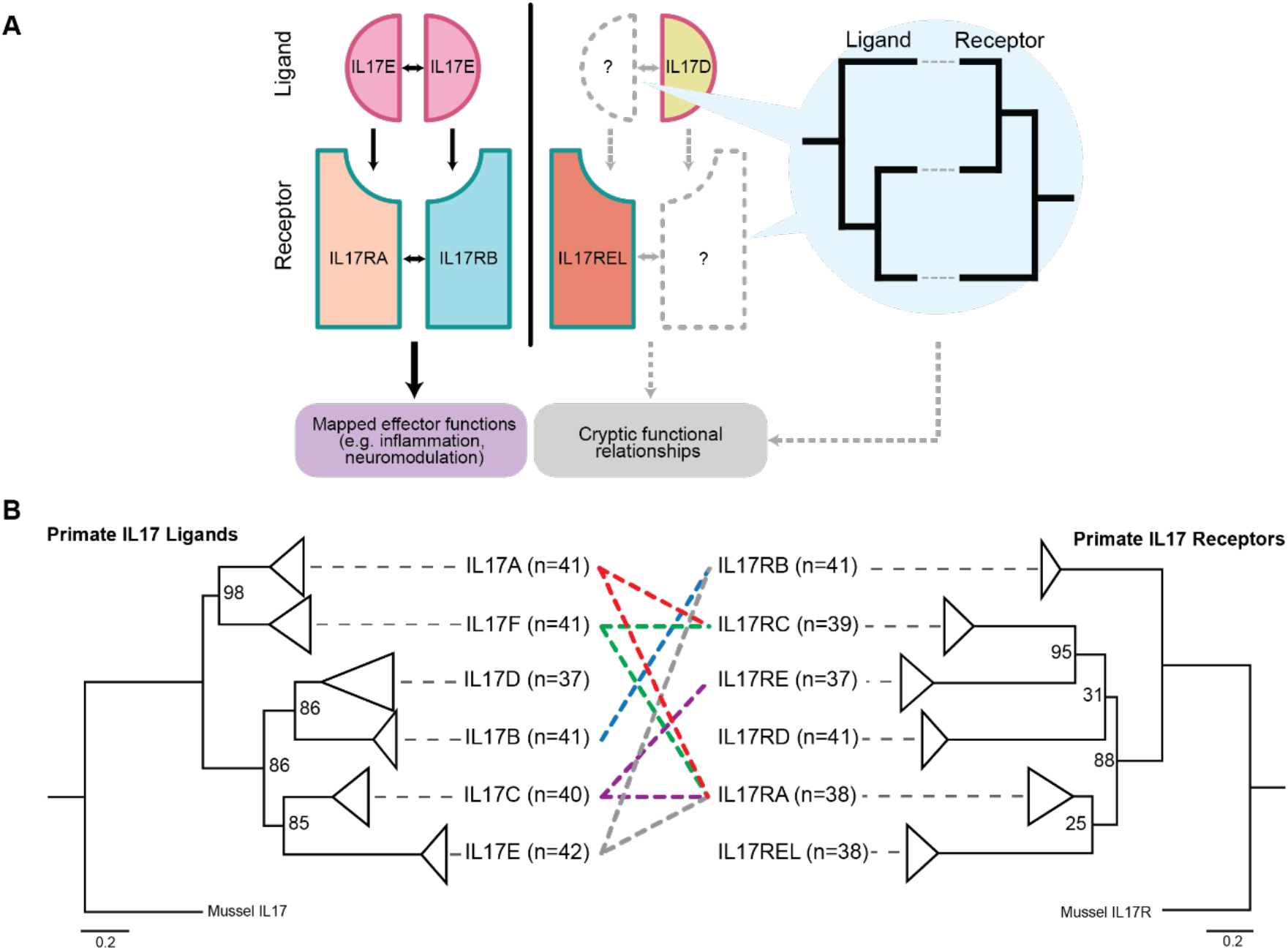
Evolutionary frameworks clarify IL-17 signaling architecture and organizing relationships. **A)** Schematic overview of the interleukin-17 (IL-17) signaling platform and the rationale for applying comparative evolutionary approaches. IL-17 ligands signal as homo- or heterodimers that engage heterodimeric IL-17 receptor (IL-17R) complexes at the cell surface (left). Canonical interactions (such as IL-17E binding to the IL-17RB/IL-17RA receptor complex) mediate defined downstream functions, including type 2 immunity and neuromodulatory signaling. However, several IL-17 subunits, including IL-17D and IL-17REL, remain poorly characterized with unresolved binding partners (middle). Phylogenetic analysis and evolutionary rate covariation (ERC) provide complementary approaches to infer co-evolving partners and uncover cryptic signaling relationships (right). **B)** Maximum likelihood phylogenetic trees of IL-17 ligands (left) and receptors (right) across ∼40 simian primate species. Distinct clades correspond to known IL-17 paralogs. Dotted lines indicate established ligand–receptor interactions. Terminal nodes are collapsed by paralog, with *n* indicating the number of orthologous simian primate sequences included per clade. Comparative topology highlights congruent ligand evolution but unexpected divergence among receptor relationships.

Consistent with these structural features, and in addition to its established role in immunity, IL-17’s role is increasingly recognized in regulating neuronal development, neuronal function, and behavioral phenotypes across multiple, divergent species (Cho, et al. 2025). The developmental outcome of this neuroimmune interface is exemplified in murine models exhibiting IL-17-dependent cortical abnormalities and behavioral changes in their offspring (Choi, et al. 2016; Shin Yim, et al. 2017; Lammert, et al. 2018; Gumusoglu, et al. 2020). Additional studies have highlighted neuromodulatory roles of IL-17 in influencing anxiety and social interaction in mice (Alves de Lima, et al. 2020; Reed, et al. 2020; Lee, Kwon, et al. 2025; Lee, Ishikawa, et al. 2025). These studies also highlight distinct IL-17R subunits specific for certain neuronal and inflammatory functions, as supported by their brain-region and cortical neuron specific expression patterns (Leonardi, et al. 2022; Lee, Kwon, et al. 2025; Lee, Ishikawa, et al. 2025). This crosstalk is not limited to vertebrates: parallel discoveries in the invertebrate nematode, *Caenorhabditis elegans*, revealed that IL-17 orthologs modulate aggregation behaviors by acting on a conserved IL-17R in neurons (Chen, et al. 2017; Godthi, et al. 2024). Beyond neurodevelopment, IL-17 dysregulation has been further associated with impaired uterine blastocyst implantation, pre-eclampsia, and other pregnancy and perinatal complications (Chavan, et al. 2017; Osborne, et al. 2019; Kwon, et al. 2022; Giangrazi, et al. 2024). Together, these findings suggest that IL-17 operates at the intersection of multiple immune, reproductive, developmental, and behavioral pathways.

Efforts to elucidate interfaces among signaling partners by solving crystal structures, such as the IL-17E– IL-17RB–IL-17RA and the IL-17A-IL-17RC-IL-17RA ternary complexes, have expanded our understanding of organizing principles. However, these studies have either excluded or modified the N-terminus disordered regions of the IL-17 ligands, which comprise nearly a third of the protein (Liu, et al. 2013; Wilson, et al. 2022). Additionally, while molecular genetics and biochemical studies have revealed key binding partners, they often failed to understand the breadth of physiological consequences. For example, IL-17D has been suggested in its role in the innate immune response, yet its cognate receptor remains unclear (Liu, et al. 2020; Huang, et al. 2021; Yuan, et al. 2025). Similarly, IL-17REL has largely remained uncharacterized since its initial discovery and is often excluded in studies characterizing IL-17 receptors (Wu, et al. 2011). Taken together, it is unclear how evolutionary pressures influence the functional diversification of the IL-17 cytokine family nor what biological features make IL-17 malleable signaling mediators.

In this study, we used evolutionary and phylogenetic methods as an orthogonal approach to better define and clarify relationships among IL-17 ligands and receptors (schematized in **Figure 1A**). To date, there have been few reports on the phylogenetic relationship between IL-17 family subunits, and available studies have been underpowered due to previously limited sequence availability. Here, we analyzed the IL-17 evolutionary histories of more than 40 primate species and discovered patterns of homolog-specific rapid evolution for both ligands and receptors, pointing to recurrent selective pressures impacting the diversification of their functions. These independent signals of positive selection have direct consequences for interpreting studies involving model organisms and for extrapolating these findings to our understanding IL-17 function in human biology. A striking example of this is found in the disordered N-terminal region of IL-17E, where we identified signatures of rapid evolution in primates, but not rodents, suggesting diverging functional properties at this interface. In addition to specific signals of recent rapid evolution, co-evolutionary rate analysis of IL-17 family members from a wide sampling of mammalian genomes highlighted a larger network of genes and signaling pathways that appear to impinge on IL-17’s multiple physiologic roles. Using these evolutionary comparisons, we propose a new framework for contextualizing the expanding and rapidly evolving functions of IL-17 and broader principles of pathway crosstalk evolution.

## Results

### Phylogenetic comparisons clarify relationships between IL-17 ligands and receptors

To better define evolutionary relationships both among IL-17 ligands and receptors and between different species, we compiled orthologous sequences for known IL-17 ligands (A–F) and receptors (RA–RE, REL) from ∼40 simian primate species (i.e. Great Apes [*Hominidae*], African/Asian monkeys [*Cercopithecidae*], and South/Central American monkeys [*Platyrrhini*]) from the TOGA ortholog database (Kirilenko, et al. 2023). Because the alphabetical naming convention of IL-17 paralogs obfuscates the relatedness of these subunits (Kawaguchi, et al. 2004) and given the dearth of species sequences used in comparative genomic studies to date, each ligand and receptor sequence sampled was manually screened for homology and subsequently used to construct maximum likelihood phylogenetic trees using PhyML (**see methods**)(Guindon, et al. 2010). Trees were rooted using IL-17 sequences found in the genome of the mussel *Mytilus galloprovincialis*, which serves as a useful outgroup for vertebrate IL-17 genes given the frequent expansion of homologs in invertebrate lineages (Saco, et al. 2021).

The resulting tree of IL-17 ligands revealed six distinct clades associated with known vertebrate paralogs IL-17A through F (**Figure 1B**). Primate tree topologies were consistent with previously established vertebrate ligand phylogenies and known genetic relationships, such as the empirical similarities between IL-17A and IL-17F. This congruence, along with robust bootstrap values of 85 or greater at terminal nodes, supports the validity of orthologous sequence detection across a wide sampling of primates. Given strong signaling associations between cognate ligand-receptor binding pairs, we hypothesized that the IL-17 receptor (IL17R) tree would follow similar patterns and mirror the evolution of ligands.

Surprisingly, patterns of receptor relatedness based on shared function were not clear in the IL-17R tree (**Figure 1B**). One example was the placement of IL-17RA; instead of clustering with any of its heterodimeric pairs IL-17RC, IL-17RB, or IL-17RE, it was placed in a distinct clade. This incongruence between ligand and receptor trees challenges assumptions about IL-17 relatedness based on function and points to the impact of genetic conflicts or environmental pressures acting independently on IL-17 ligands and receptors.

In addition, we found evidence clarifying the phylogenetic relationships of orphaned receptors IL-17RD and IL-17REL (RE-like) in comparison to other subunits. Notably, IL-17REL was initially named for its perceived similarity to IL-17RE, but our analysis suggests it is more closely related to IL-17RA in primates (**Figure 1B**). This revised placement reveals a red herring in the naming of IL-17REL, and we hypothesize that its effector functions may be more closely aligned with those of IL-17RA than previously appreciated. Indeed, a recent study indicates that IL-17REL may function as a decoy receptor, suppressing IL-17A, IL-17F, and IL-17C activity by directly binding to these ligands, likely through competition with IL-17RA (Li, et al. 2026). This finding further validates the phylogenetic signals of IL-17 evolution.

Next, we made phylogenetic comparisons of IL-17 subunits between animal lineages, with a focus on rodents given the prevalence of studies of IL-17 based on murine models. Using the same pipeline described above, we created phylogenetic trees for IL-17 ligands and receptors by comparing sequences from ∼40 rodent species. While IL-17 cytokine family subunits are generally assumed to be conserved across vertebrates, gene tree comparisons between primates and rodents revealed marked contrasts (**Figure 2A, B**). Like primates, the rodent ligand tree resolved into six distinct clades for each IL-17 paralog. However, the placement of IL-17E between primates and rodents was notably different, with IL-17E at the root of all ligands in rodents compared to a more recent branchpoint inferred with IL-17C in primates. This suggests that IL-17E is impacted by distinct evolutionary pressures and raises the possibility that specified functions have evolved for IL-17E in each animal lineage.

**Figure 2.**
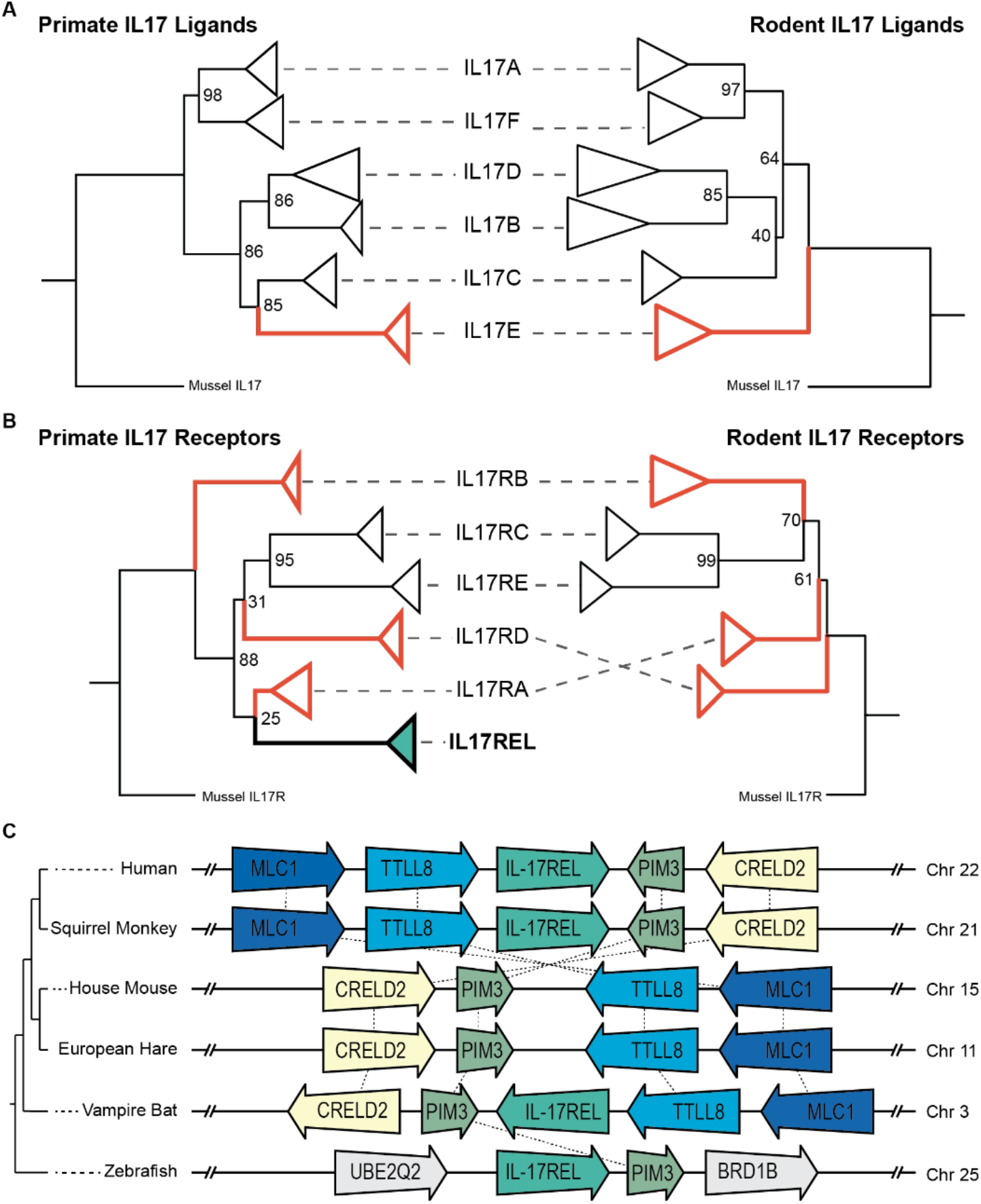
Lineage-specific diversification defines IL-17 ligand and receptor evolutionary histories. **A–B)** Tanglegram comparisons of IL-17 ligand (**A**) and receptor (**B**) phylogenies between primates (left) and rodents (right) reveal tree topology indicative of lineage-specific evolutionary trajectories. Red branches/lineages indicate incongruent branching patterns between rodents and primates, emphasizing divergence in ortholog relationships across mammals. Bootstrap support values are shown for major nodes. **C)**Schematic of genomic synteny analysis of the IL-17REL locus across representative vertebrate species referenced in this study. Conserved flanking regions provide genomic context of this region while the IL-17REL coding sequence is selectively pseudogenized in rodents and lagomorphs. This pattern supports lineage-specific gene loss as a mechanism shaping IL-17 pathway composition.

In addition, receptor groupings among rodents exhibited multiple rearrangements compared to phylogenetic patterns observed for primates. This volatility in receptor evolution might explain how IL-17 evolution studies to date have focused exclusively on ligand diversification given more defined points for comparison. However, our analysis highlights remarkable diversification of IL-17Rs, emphasizing an understudied trove of possible species-specific IL-17 pathway usage. For example, we confirmed that the IL-17REL loci has been pseudogenized and degraded in rodents and lagomorphs, as evidenced by its degradation in the syntenic region shared from mammals to teleost fishes (**Figure 2C**). Taken together, our results suggest IL-17 evolution is much more fluid than previously anticipated, as distinct ligand and receptor phylogenies indicate unique functional diversification between animal lineages.

### Diversification of IL-17 is marked by recurrent rapid evolution

Given the incongruent IL-17 topologies and drastic variability between mammalian clades, we sought to test the hypothesis that these changes were a result of recurrent positive selection acting at ligand-receptor interfaces, a pattern observed for many genes encoding immune signaling functions (Bustamante, et al. 2005; Nielsen, et al. 2005; Shultz and Sackton 2019). Evidence of rapidly evolving cytokines could point to genetic or environmental pressures underlying the neofunctionalization of IL-17 and/or pinpoint specific residues that disproportionately influence IL-17 signaling. We aligned and manually trimmed orthologous IL-17 ligands and receptors and tested for signatures of positive selection using several codon-based selection algorithms including PAML (Yang 2007) and FUBAR (Murrell, et al. 2013), MEME (Murrell, et al. 2012), and BUSTED (Murrell, et al. 2015) from the HyPhy suite (Pond, et al. 2004). These selection models provide complementary methods to detect evidence of both episodic and pervasive selection across phylogenies.

Using these site models, we identified nine of the 12 primate IL-17 subunits evolving under positive selection in at least one selection model (**Figure 3A**). Mapping positively evolving subunits onto the IL-17 phylogenetic tree revealed that evolutionarily grouped subunits followed similar patterns of positive selection. Among the pervasive signals of positive selection, the ligand IL-17E and its receptor IL-17RB and IL-17REL, exceeded significance thresholds for all selection models tested; given IL-17E and IL-17RB’s emerging roles in both neuromodulatory function and recently identified anti-inflammatory function of IL-17REL, we focused further evolutionary analyses on this triad (Lee, Ishikawa, et al. 2025; Li, et al. 2026).

**Figure 3.**
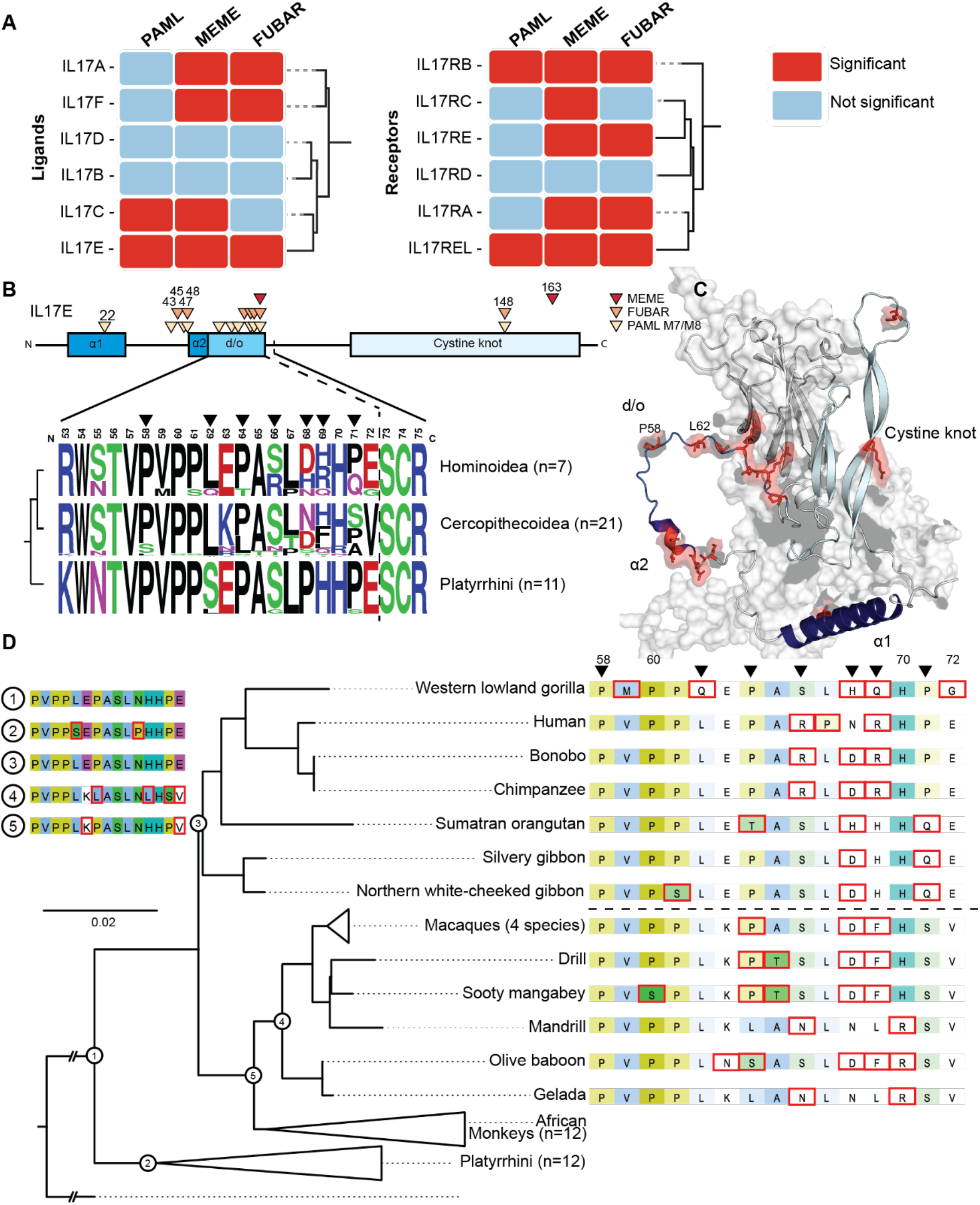
Positive selection drive rapid diversification of primate IL-17 subunits. **A)** Summary of positive selection analyses across simian primate IL-17 ligands and receptors supported by PAML (BEB > 0.99), MEME (p < 0.05), and FUBAR (posterior probability > 0.95). Multiple subunits exhibit signatures of episodic and/or pervasive positive selection, indicating recurrent adaptive pressures. **B)** Schematic of the IL-17E domain architecture highlighting residues under positive selection. Rapidly evolving sites cluster within the N-terminal disordered domain, whereas the conserved cysteine-knot receptor binding region remains largely constrained. **C)** Structural model of the IL-17E:IL-17RB monomeric complex, as predicted by AlphaFold. The N-terminal disordered region and α-helical elements are shown, with positively selected residues highlighted in red, illustrating spatial segregation between conserved structural interfaces and rapidly evolving regions. **D)** Partially collapsed phylogenetic tree of IL-17E across simian primates. Residues outlined in red indicate derived amino acid substitutions relative to the most recent reconstructed ancestral state, demonstrating lineage-specific diversification concentrated in the N-terminal domain.

Curiously, despite the strong signals of positive selection in primates, when we compared primate IL-17 subunits to other mammalian clades, we found that, when normalized for tree length, the same orthologs in rodents and bats have different signatures of positive selection (**Supplemental Figure 1**). For example, the strong signal for selection in IL-17E was abolished in rodents and alternate residues were rapidly evolving in bats. This prompted us to investigate these patterns in more depth.

### Neuronally expressed IL-17E residues in its N-terminus disordered domain are rapidly evolving

As judged by three episodic diversification models, the strongest signatures of positive selection were evident in IL-17E ligand and IL17RB and IL17REL receptors. The IL-17E:IL-17RB interface was a striking find, given the recent reports that this cognate ligand-receptor pair is highly expressed in neuronal tissues and is associated with previously reported neurodevelopmental phenotypes (Lee, Kwon, et al. 2025; Lee, Ishikawa, et al. 2025). Additionally, IL-17E has been implicated in mounting immune responses against parasitic infection and type 2 immunity (Reynolds, et al. 2010; Howitt, et al. 2016; von Moltke, et al. 2016; Schneider, et al. 2019), highlighting the IL-17E:IL-17RB interface as an apt model to investigate the evolution of crosstalk between disparate physiologic processes.

Given diverse biological relevance of these signals, we sought to identify specific residues evolving under positive selection in IL-17E and discovered that the signal was focused on a stretch of 15 amino acids in the N-terminus disordered domain (**Figure 3B**). In sharp contrast, the cysteine knot receptor-binding domain directly downstream—which binds to its cognate receptor, IL-17RB—is highly conserved (**Figure 3C**). Little is known about the ∼70 amino acid N-terminal region contained in all IL-17 ligands; given the instability of disordered domains, structural studies and protein purifications of IL-17 ligands have often omitted this rapidly evolving region.

Thus, to better understand the pervasive changes occurring to this N-terminal disordered region, we used ancestral sequence reconstruction (ASR) to assess patterns of mutations that arose during the evolution of primates **(Figure 3D)**. ASR revealed that IL-17E is highly conserved in South/Central American monkeys. Conversely, the evolutionary signature is heavily influenced by sequence heterogeneity in apes and Asian/African monkeys, two clades separated by ∼28 million years (**Figure 3D**). Thus, adaptive strategies to evolutionary selective pressures on the N-terminus region and the IL-17E:IL-17RB interface appear to be acting independently, even between closely related clades.

Ancestral reconstructions also revealed multiple instances of convergent acquisition of amino acids. In several of these instances, the convergent acquisitions span different amino acid side chain groupings, such as independently derived histidines and aspartic acids from the ancestral asparagine on residue 68 (**Figure 3D**). Additionally, across the 15-amino acid domain, there have been many substitutions to or from prolines. Furthermore, ASR revealed a primate-specific eight-amino acid C-terminal deletion shared among Asian/African monkeys and great apes (**Figure 3D, Supplemental Figure 2**), suggesting a derived adaptation that may influence receptor binding or downstream signaling. Taken together, the widespread sequence heterogeneity, the diversity of amino acid substitutions, and the clade-specific loss of coding region – all within a condensed evolutionary timeframe – point to a region of intense evolutionary volatility within IL-17E.

### Covariation analysis underscores a complex evolutionary history of IL-17 factors

Maximum likelihood methods are powerful tools to identify rapidly evolving genes and residues with likely functional consequences. However, these approaches are agnostic to the broader context of biologic networks. To integrate our understanding of how positively selected subunits like IL-17E and IL17RB are impacting related pathways, we used evolutionary rate covariation (ERC) to compare the relative differences in evolutionary rates of IL-17 family members. ERC captures the tendency of functionally linked genes to co-vary at similar mutation rates allowing for identification of hidden genetic relationships missed in *in vitro* settings (Clark, et al. 2012; Little, et al. 2025). Importantly, ERC is a strong indicator of co-functional associations, and not binding, which makes it a complimentary foil to binding and pull-down studies which may miss subtle interactions and co-factors(Little, et al. 2024).

Among all IL-17 subunits, IL-17E and its receptor IL-17RB had a particularly high ERC value across mammals (**Figure 4A**). This coevolutionary signal is notable given that both IL-17E and IL-17RB have multiple binding subunits; promiscuity in binding partners weakens coevolutionary signals, likely due to compensatory adaptations being distributed across multiple parameters. For example, IL-17A, despite having many known strong interaction partners, showed significantly weaker ERC values for its cognate receptors (such as IL-17RA and IL-17RC and for heterodimer ligand IL-17F; **Figure 4A**); in this instance, the ERC score below our threshold of significance may imply functional flexibility of IL-17A. However, the IL-17E:IL-17RB pair appears to defy this trend. Despite evidence of multiple binding pairs (akin to IL-17A), their strong ERC signal, together with evidence of lineage-specific positive selection denoted above, suggests that this pair is under distinct evolutionary pressures, such as emerging specialization beyond their canonical immune roles.

**Figure 4.**
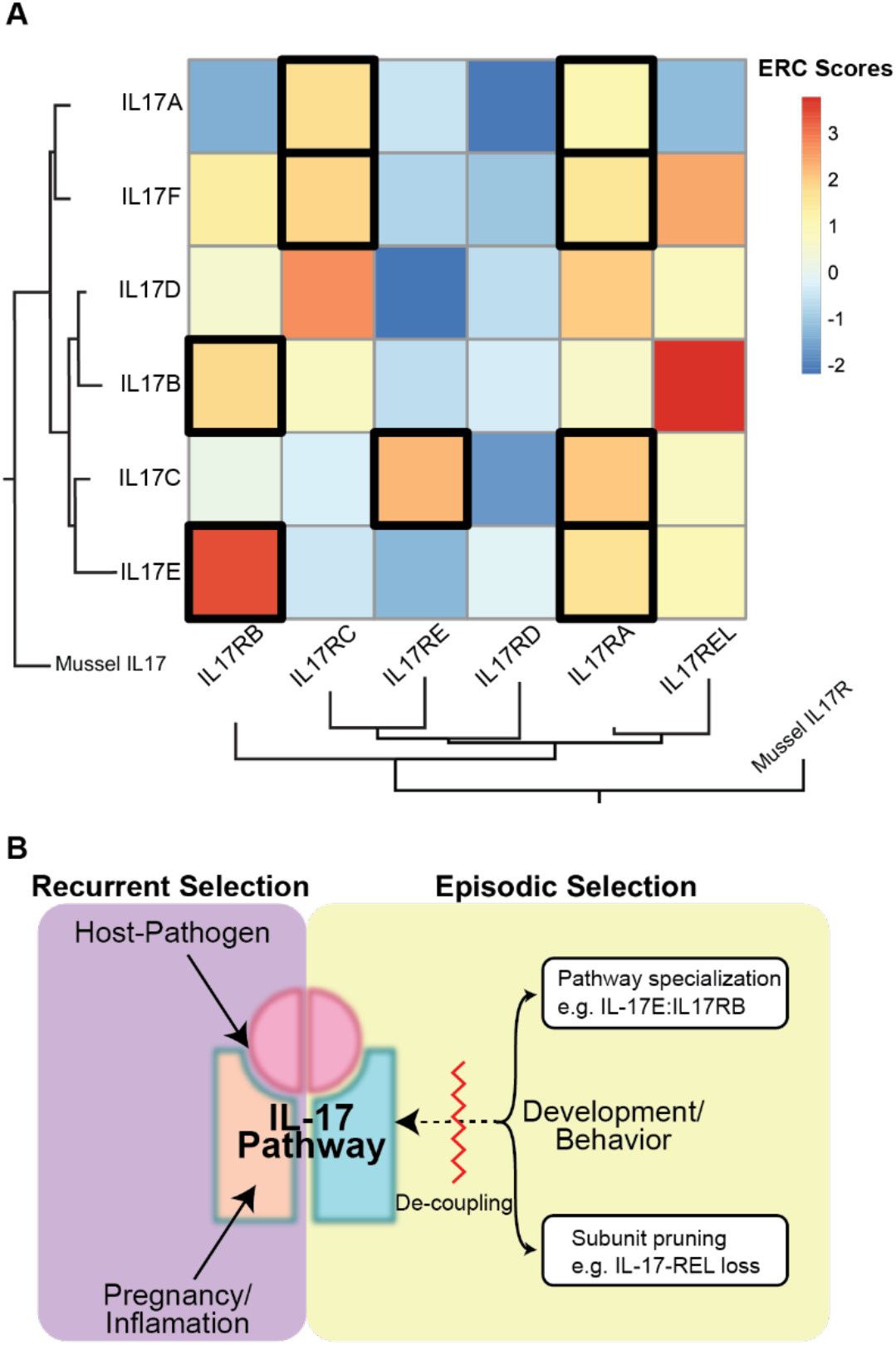
Co-evolutionary analysis underscore the complex networks influencing IL-17 diversification. **A)** Heatmap of evolutionary rate covariation (ERC) scores among IL-17 ligands and receptors across mammals. Higher ERC values indicate stronger co-evolutionary relationships. Black boxes denote known ligand–receptor pairs, while additional high-scoring interactions suggest previously unrecognized functional associations (e.g. IL-17D:IL-17RC and IL-17B:IL-17REL). **B)** Conceptual model for the evolution of IL-17 signaling under competing selective pressures. Recurrent pressures (e.g., host–pathogen interactions) drive rapid diversification and dominate pathway evolution, whereas more constant pressures (e.g., developmental or life-history constraints) are resolved through two mechanisms: (1) Pathway specialization, where specific paralogs (e.g., IL-17E/IL-17RB) diverge toward dedicated non-immune roles such as neurodevelopmental signaling; and (2) Subunit pruning, where genes with conflicting functional demands (e.g., IL-17REL in rodents) are lost. This framework provides a generalizable model for understanding the evolution of signaling pathway crosstalk under competing biological constraints.

Given these coevolutionary signals, one intriguing hypothesis is that IL-17E and IL-17RB are becoming increasingly specialized for neuron- and neurodevelopment-specific functions, where precise ligand-receptor interactions may be more stringently regulated. This idea is further reinforced by observations that IL-17E is more strongly co-evolving with other neurodevelopmental genes, such as *Noggin*, than members of inflammatory pathways (**Supplemental Table 1**). In this view, the evolutionary pressure to maintain functional fidelity in neural signaling may outweigh the broader flexibility typically seen in immune signaling. Therefore, strong patterns of co-evolution between IL-17E and IL-17RB, may suggest a shift toward tissue-specific roles, potentially reflecting either an adaptive transition to neurodevelopmental regulation or away from immune signaling.

ERC analysis also revealed a broader network of hidden evolutionary relationships across not only IL-17 genes but also with genes in non-immune pathways (**Figure 4A, Supplemental Table 1, Supplemental Table 2**). Strikingly, the strongest ERC signal between IL-17 subunits corresponds to IL-17REL and IL-17B. While a recent report indicated that IL-17REL primarily antagonizes the function of IL-17A, IL-17F, and IL-17C, not IL-17B (Li, et al. 2026),our ERC analyze suggest a potential functional interaction between IL-17REL and IL-17B. One possibility is that IL-17B must need to form a complex with its cognate receptor IL-17RB before engaging IL-17REL, a hypothesis that will require future investigation.

We also identified strong co-evolutionary signatures between the ligand IL-17D and the receptor IL-17RC. Very little is known about IL-17D (Liu, et al. 2020; Huang, et al. 2021; Yuan, et al. 2025), but these data suggest the presence of a novel cryptic signaling relationship between these two subunits, whose functions have been elusive using conventional assays. These data, thus, highlight an intriguing hypothesis to test if IL-17D functionally interacts with IL-17RC and its interacting partner IL-17RA.

## Discussion

Fidelity of interleukin signaling is essential for marshalling effective host defenses against pathogens. However, immune-dedicated ligands and receptors commonly diversify and encode additional signaling processes, repurposing existing signaling relays for tissue-specific and/or non-immune functions (McGeachy, et al. 2019; Ferro, et al. 2021; Kim, et al. 2024). This biological complexity has the potential to entangle overlapping processes and cloud our ability to reconcile conflicting functions when multiple biological outcomes are orchestrated through shared ligands and receptors.

For IL-17, mounting evidence shows that this signaling platform participates in a wide network of discrete physiologic processes (i.e. host-pathogen, maternal-fetal, neurobehavioral, and developmental interfaces (Choi, et al. 2016; Chavan, et al. 2017; Chen, et al. 2017; Osborne, et al. 2019; Reed, et al. 2020; Kwon, et al. 2022; Giangrazi, et al. 2024; Godthi, et al. 2024; Cho, et al. 2025; Lee, Kwon, et al. 2025; Lee, Ishikawa, et al. 2025)) each distinctly impacted by its respective environmental cues. Thus, this study took a comparative evolutionary approach to not only clarify the selective pressures driving diversification of the IL-17 pathway, but to study IL-17 as a model for how complex pathway crosstalk is influenced by competing environmental cues. Our findings advance the IL-17 system as a heuristic for understanding how a wide range of both recurrent and more constant evolutionary pressures might diversify this – and other – immune signaling pathways such as chemokines and other innate and adaptive immune receptors (**Figure 4B**).

We first established phylogenetic relationships between IL-17 ligands and receptors (**Figure 1B**) and between different species (**Figure 2**) and found patterns congruent with ongoing, recurrent adaptation. Indeed, similar to many immune pathways, we observed pervasive signals of rapid evolution in IL-17 ligands and receptors consistent with Red Queen dynamics driven by recurrent genetic conflicts with infectious agents (Nielsen, et al. 2005; Shultz and Sackton 2019) (**Figure 3**). These findings suggest that IL-17 variants are selected to regain or enhance immune responses that counter ongoing adaptive changes in pathogens at host protein interfaces. Specifically, analysis of recurrent positive selection points to an underappreciated disordered loop in IL17E at the interface with IL17RB (**Figure 3B**). These patterns of evolutionary change are consistent with the impact of specific pathogen-encoded proteins that might disrupt or alter functions at this interaction site between IL17E and IL17RB (Howitt, et al. 2016; von Moltke, et al. 2016; Schneider, et al. 2019). However, IL-17E and IL-17RB’s crosstalk complicates this paradigm; in addition to considering the consequences of pathogen-driven evolution, growing evidence places the IL-17 pathway in regulating multiple aspects of neural activity and function, particularly in the contexts of social interaction, anxiety, and memory. In contrast to genetic conflicts resulting in recurrent adaptations, the impact of more consistent biological cues, such as behavior, life-history traits, and developmental processes, often emerges as more rare adaptive events. However, it has been unclear how these pressures compete with immune processes and ultimately influence crosstalk.

We propose a model in which IL-17 signaling has been impacted by competing demands from immune interfaces which outweigh repercussions for archetypal development and behavior related signaling (**Figure 4B**). Our analysis suggests a model where selection acts on gene duplication and gene loss events that facilitate IL-17 neuroimmune crosstalk, while delineating development- and behavior-associated functions. Several lines of evidence support the hypothesis of neurobehavioral pathway specialization. Recent work showed that IL-17E and the receptor IL-17RB are expressed in neuronal tissue and directly modulate social behaviors (Lee, Ishikawa, et al. 2025). This unique role is echoed in our phylogenetic reconstruction placing IL-17RB in a clade distinct from other related receptors (**Figure 1B**). Additionally, the placement of IL-17E varies drastically between species, in contrast to conservation of other IL-17 ligand paralogs (**Figure 2A**). Furthermore, in primates, we uncovered strong co-evolutionary signatures as well as signatures of positive selection between IL-17E and IL-17RB (**Figures 3A and 4A**).

We interpret these data as evidence for pathway specialization, where paralogs have adapted to support a distinct IL-17 signaling axis involved in development and/or behavior. Further studies of lineage-specific IL-17 gains or losses may help elucidate the impact of subunit pruning. Taken together, we believe future experiments should prioritize the N-terminal disordered domain of IL-17E and the standalone potential in the neuronal IL-17E and IL-17RB signaling axis on behavior and the tradeoffs of losing this axis when mounting immune responses to pathogens.

Additionally, our analysis suggests new roles for understudied IL-17 subunits, such as IL-17REL, including candidate binding partners. Relatively few studies have addressed the role of IL-17REL in immune signaling and/or other related functions mediated by IL-17 paralogs (Li, et al. 2026). Evidence from phylogenetic analysis in a sampling of primates raises the intriguing possibility of a role for IL-17REL as a neurodevelopmental and behavioral modulator of the IL-17 neuroimmune pathway. Its phylogenetic placement with IL-17RA, highly correlated evolutionary rates between IL-17REL and IL-17B, and strong signatures of positive selection across the three neuronally expressed subunits lend support to this idea (**Figure 4A**). Future work testing candidate interactions with IL-17REL will provide a new understanding of the scope and potential of IL-17 functions in primates.

Of particular interest, we describe the genetic loss of IL-17REL in rodents and lagomorphs (**Figure 2**). Rather than retaining and repurposing the pathway for multiple, independent functions, gene loss is another means to mediate a range of overlapping and/or conflicting neuromodulatory IL-17 functions. Similar patterns of adaptive loss have been documented elsewhere, such as in the OAS1/RNase L and APOBEC anti-viral effector families, where gene loss events might resolve harmful pleiotropy and genetic conflicts (OhAinle, et al. 2006; Carey, et al. 2019). Curiously, in the species considered for this study, we have not observed evidence for concurrent pathway specialization *and* loss of subunits; this observation supports the hypothesis that species are independently exploring evolutionary strategies to refine IL-17’s diverse biological functions (**Figure 4B**).

Importantly, the loss of IL-17REL in rodents highlights the limitations of relying solely on mouse models to reveal the full scope of *in vivo* IL-17 functions. In this context, zebrafish represent an intriguing alternative model, given the presence of syntenic IL-17REL and other prominent players in the IL-17 pathway, especially in light of established behavioral paradigms to test sociability phenotypes (Kono, et al. 2011; Wu, et al. 2011; Nelson and Granato 2022; González-Fernández, et al. 2024). Alternatively, it would be useful to develop a rodent model that expresses IL-17REL under physiologically relevant regulatory control (Li, et al. 2026). However, interpretation of such models should be contextualized by our findings that rodent and lagomorph IL-17 complexes have independently evolved and specialized without IL-17REL for approximately 90 million years (Kumar, et al. 2017).

Another factor to consider in IL-17 evolution is the recurrent selective pressure of IL-17 in pregnancy and maternal-fetal conflicts (**Figure 4B**). Both sexual conflict and host–pathogen arms races, can result in similar signals of rapid evolution, a key consideration for advancing hypotheses and pursuing future tests of the functional impacts of rapid evolution at these interfaces involving IL-17 (Daugherty and Malik 2012).

By embracing the complexity of IL-17’s tangled evolutionary past, we are better positioned to understand not only its immunologic roles, but also its wide-ranging contributions from neurological homeostasis and diseases to altered behavioral outcomes. Our findings suggest that future research on the IL-17 system would benefit from adopting a broader framework for interpreting IL-17 functions, beyond canonical immune roles. Especially considering the clade-specific diversification of subunits, such as IL-17E, interpretation of data from experiments in animal models in the context of human biology should be weighed carefully. By placing the diversity of IL-17 functions in an evolutionary framework, we provide a platform for better understanding how biological forces are shaping the functions and adaptations of this key pathway, and how analogous forces influence the evolution of other signaling networks. Although the evolutionary hypotheses proposed here remain to be experimentally tested, this work helps clarify and prioritize research questions and directions for the field.

**Supplemental Figure 1.**
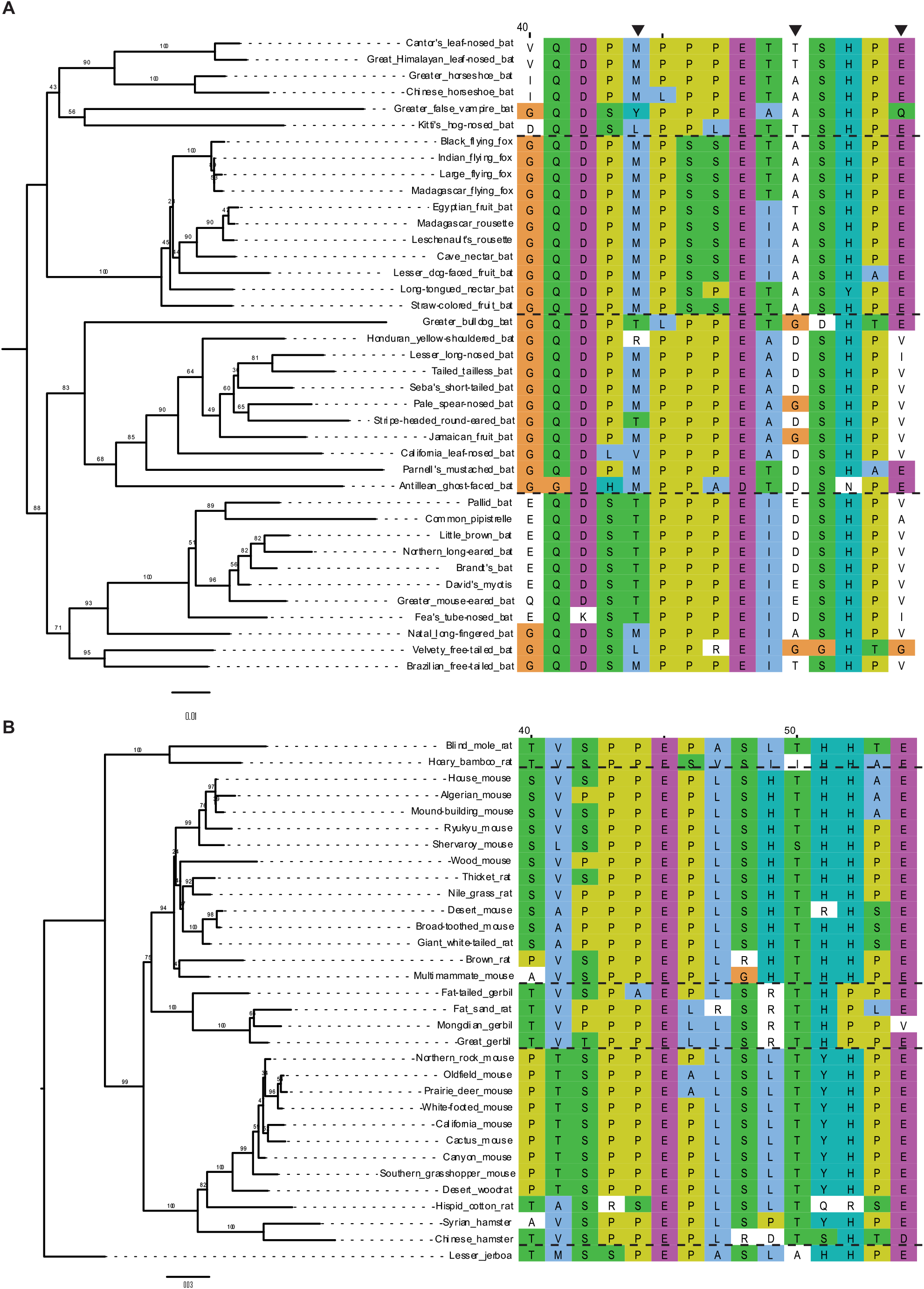
Maximum likelihood gene tree of A) bat and B) rodent IL-17E with alignment of 15 amino acid N-terminal disordered region under pervasive positive selection in primates. Black arrows represent residues under rapid evolution as determined by PAML M7-M8 model. PAML did not register rodent IL-17E as evolving under selection.

**Supplemental Figure 2.**
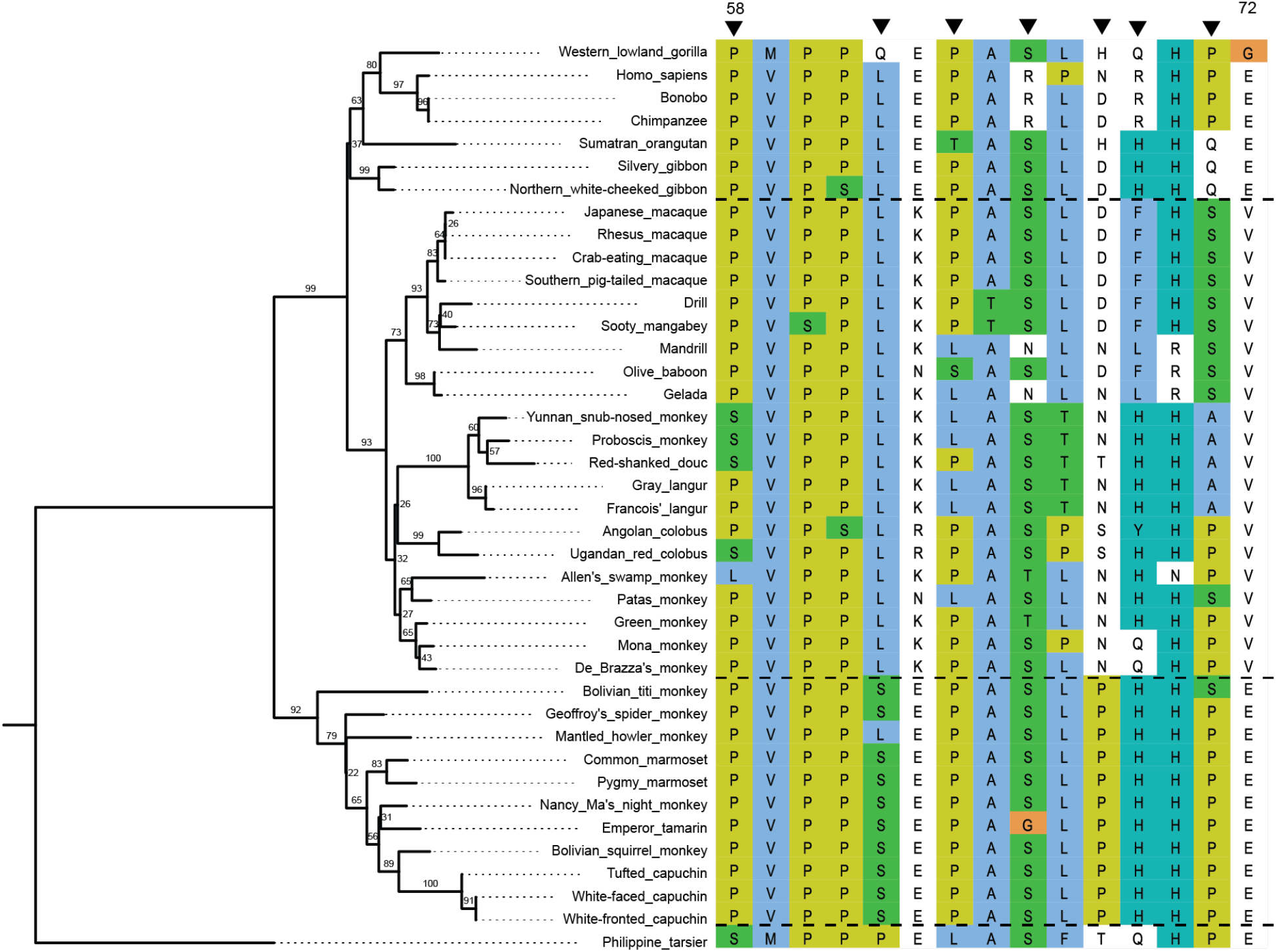
Uncollapsed maximum likelihood gene tree of primate IL-17E orthologs used in this study. Alignment window shows the N-terminus disordered domain and black arrows indicate rapidly evolving residues as determined by PAML (BEB > 0.99), MEME (p < 0.05), and/or FUBAR (posterior probability > 0.95).

## Materials and Methods

### IL-17 Sequences and alignments

Orthologous gene sequences for IL-17 ligands and receptors were obtained from the Zoonomia Project TOGA ortholog database, using the human genome assembly hg37 as the reference build (Kirilenko, et al. 2023). All available sequences for Simiiformes (for simian primate clades), Muroidea (for rodent clades), and Chiroptera (for bat clades) on Zoonomia were compiled for subsequent analysis. The TOGA database uses machine learning algorithms to infer orthologous gene loci based on a query sequence. Although this yields a high-throughput approach of identifying orthologs based on both sequence identity and location, there are instances where this algorithm falls short. Thus, we found that not all simian primate species had complete, annotated IL-17 orthologs with some missing entirely from queries and multiple instances where sequences were degraded or missing key identifiers, such as start and stop codons. Where possible, we substituted these orthologs with publicly available sequences in the NCBI (Sayers, et al. 2024) or ENSEMBL (Dyer, et al. 2024) gene databases. In addition, we validated one-to-one ortholog comparisons by manually checking all genes for synteny in the final curated sequence compilation.

Multiple codon alignments were generated for IL-17 ligand and receptor subunit using Clustal Omega build 1.2.2 with default parameters (Sievers, et al. 2011; Madeira, et al. 2024). We inspected subsequent alignments for poorly annotated or low confidence regions, ambiguous codons, or gaps in the sequence, and omitted these from further analysis. When applicable, we visualize amino acid conservation of aligned domains using sequence logo plots generated by the Berkeley WebLogo server (Crooks, et al. 2004).

### Phylogenetic and synteny analyses

We established phylogenetic trees with IQ-TREE version 1.6.12 using ModelFinder and PhyML build 3.3.20180621 using both JC69 and HKY85 substitution models with 100 bootstrap replicates to assess node support (Guindon, et al. 2010). All models consistently yielded congruent topologies, thus we applied PhyML and JC69 thereafter for consistency and to optimize computational burden across all IL-17 orthologs. Trees were visualized and annotated in FigTree v1.4.4 and longer branch lengths were manually truncated to preserve branchlength scales (Rambaut and Drummond 2012). For direct comparison of alternative topologies, we used Dendroscope V3_8_10 to generate tanglegrams and infer branch and clade rearrangements (Huson, et al. 2007). As mentioned above, we confirmed genomic synteny for all sequences to ensure ortholog comparisons and to evaluate the genomic context of target loci. We identified orthologous flanking genes using NIH NCBI gene database (Sayers, et al. 2024) in representative species, and synteny maps were compared across genomes to establish conserved gene order.

### Ancestral sequence reconstruction

To calculate ancestral nodes, we ran IL-17 sequences through FireProt-ASR v.2.0 webserver as outlined in Musil, et al. 2020 (Musil, et al. 2020). Input alignments were curated as described above in Sequences and alignments. Resulting sequences were superimposed on species trees referenced from TimeTree (Kumar, et al. 2017).

### Selection analysis and estimation of substitution rates

We conducted codon-based selection analyses on IL-17 alignments to identify rapidly evolving residues with high likelihood of functional consequences. Multiple, complementary approaches were employed for this study: PAML M7 vs M8 and M8 vs M8a for branch-site likelihood ratio tests, and MEME and FUBAR (HyPhy package) for site-level selection inference (Pond, et al. 2004; Yang 2007; Murrell, et al. 2012; Murrell, et al. 2013; Murrell, et al. 2015). Scripts and parameters for all selection analyses are included below. Statistical significance was assessed as such: PAML (BEB > 0.99), MEME (p < 0.05), and FUBAR (posterior probability > 0.95).

### Evolutionary rate covariation (ERC) analysis

ERC scores and matrices were obtained using networks and evolutionary distances established by Nathan Clark’s group and subsequent protocols (Clark, et al. 2012; Little, et al. 2024; Little, et al. 2025). Pairwise evolutionary rate covariation values were computed for all possible IL-17 ligand and receptor pairs and resulting correlation matrices were visualized using pheatmap v.1.0.13 (Kolde 2019).

### Structural Modeling and Visualization

Predicted protein structures were generated with sequences obtained from Zoonomia or NCBI using AlphaFold3 (Abramson, et al. 2024). Structural models were visualized and annotated using PyMOL v.3.0.2 to highlight conserved residues and putative functional sites identified from sequence and evolutionary analyses.

## Acknowledgements

This work was supported by the University of Utah Genetics Training Program T32GM141848, F30AI191743 to S.C. and NIH grant R35GM134936 to N.C.E. We thank members of the Elde Lab for their feedback and support on this project. We thank Jordan Little and Nathan Clark for their input and advice on evolutionary rate covariation analyses in this study. We thank Gillian Stanfield, David Grunwald, and T32 Utah Genetics Training Program Fellows for helpful discussions.

## Data Availability Statement

All sequence data used in this study were obtained from publicly available databases: TOGA ortholog database (Kirilenko, et al. 2023), NCBI (Sayers, et al. 2024), and ENSEMBL (Dyer, et al. 2024). Raw sequences, alignments, expanded phylogenetic trees newick files, dn/ds and selection analysis codes/outputs, and complete ERC comparisons are available in supplemental files.

## References

1. Abramson J, Adler J, Dunger J, Evans R, Green T, Pritzel A, Ronneberger O, Willmore L, Ballard AJ, Bambrick J, et al. 2024. Accurate structure prediction of biomolecular interactions with AlphaFold 3. Nature 630:493–500.

2. Alves de Lima K, Rustenhoven J, Da Mesquita S, Wall M, Salvador AF, Smirnov I, Martelossi Cebinelli G, Mamuladze T, Baker W, Papadopoulos Z, et al. 2020. Meningeal γδ T cells regulate anxiety-like behavior via IL-17a signaling in neurons. Nature Immunology 21:1421–1429.

3. Brocker C, Thompson D, Matsumoto A, Nebert DW, Vasiliou V. 2010. Evolutionary divergence and functions of the human interleukin (IL) gene family. Human Genomics 5:30.

4. Bustamante CD, Fledel-Alon A, Williamson S, Nielsen R, Todd Hubisz M, Glanowski S, Tanenbaum DM, White TJ, Sninsky JJ, Hernandez RD, et al. 2005. Natural selection on protein-coding genes in the human genome. Nature 437:1153–1157.

5. Carey CM, Govande AA, Cooper JM, Hartley MK, Kranzusch PJ, Elde NC. 2019. Recurrent Loss-of-Function Mutations Reveal Costs to OAS1 Antiviral Activity in Primates. Cell Host & Microbe 25:336-343.e334.

6. Chavan AR, Griffith OW, Wagner GP. 2017. The inflammation paradox in the evolution of mammalian pregnancy: turning a foe into a friend. Current Opinion in Genetics & Development 47:24–32.

7. Chen C, Itakura E, Nelson GM, Sheng M, Laurent P, Fenk LA, Butcher RA, Hegde RS, de Bono M. 2017. IL-17 is a neuromodulator of Caenorhabditis elegans sensory responses. Nature 542:43–48.

8. Chen S, Fan H, Ran C, Hong Y, Feng H, Yue Z, Zhang H, Pontarotti P, Xu A, Huang S. 2024. The IL-17 pathway intertwines with neurotrophin and TLR/IL-1R pathways since its domain shuffling origin. Proceedings of the National Academy of Sciences 121:e2400903121.

9. Cho DH, Huh JR, Choi GB. 2025. Neuromodulation by the immune system: implications for brain-directed immunotherapy. Current Opinion in Immunology 95:102568.

10. Choi GB, Yim YS, Wong H, Kim S, Kim H, Kim SV, Hoeffer CA, Littman DR, Huh JR. 2016. The maternal interleukin-17a pathway in mice promotes autism-like phenotypes in offspring. Science 351:933–939.

11. Clark NL, Alani E, Aquadro CF. 2012. Evolutionary rate covariation reveals shared functionality and coexpression of genes. Genome research 22:714–720.

12. Crooks GE, Hon G, Chandonia JM, Brenner SE. 2004. WebLogo: a sequence logo generator. Genome Res 14:1188–1190.

13. Daly AE, Chang AB, Purbey PK, Williams KJ, Li S, Redelings BD, Yeh G, Wu Y, Pope SD, Venkatesh B, et al. 2025. Stepwise neofunctionalization of the NF-κB family member Rel during vertebrate evolution. Nature Immunology 26:760–774.

14. Daugherty MD, Malik HS. 2012. Rules of Engagement: Molecular Insights from Host-Virus Arms Races. Annual Review of Genetics 46:677–700.

15. Ferro A, Auguste YSS, Cheadle L. 2021. Microglia, Cytokines, and Neural Activity: Unexpected Interactions in Brain Development and Function. Frontiers in Immunology Volume 12-2021.

16. Gaffen SL. 2009. Structure and signalling in the IL-17 receptor family. Nature Reviews Immunology 9:556–567.

17. Giangrazi F, Buffa D, Lloyd AT, Redmond AK, Glover LE, O’Farrelly C. 2024. Evolutionary Analysis of the Mammalian IL-17 Cytokine Family Suggests Conserved Roles in Female Fertility. American Journal of Reproductive Immunology 92:e13907.

18. Godthi A, Min S, Das S, Cruz-Corchado J, Deonarine A, Misel-Wuchter K, Issuree PD, Prahlad V. 2024. Neuronal IL-17 controls <i>Caenorhabditis elegans</i> developmental diapause through CEP-1/p53. Proceedings of the National Academy of Sciences 121:e2315248121.

19. González-Fernández C, García-Álvarez MA, Cuesta A. 2024. Identification and functional characterization of fish IL-17 receptors suggest important roles in the response to nodavirus infection. Marine Life Science & Technology 6:252–265.

20. Guindon S, Dufayard J-F, Lefort V, Anisimova M, Hordijk W, Gascuel O. 2010. New Algorithms and Methods to Estimate Maximum-Likelihood Phylogenies: Assessing the Performance of PhyML 3.0. Systematic Biology 59:307–321.

21. Gumusoglu SB, Hing BWQ, Chilukuri ASS, Dewitt JJ, Scroggins SM, Stevens HE. 2020. Chronic maternal interleukin-17 and autism-related cortical gene expression, neurobiology, and behavior. Neuropsychopharmacology 45:1008–1017.

22. Howitt MR, Lavoie S, Michaud M, Blum AM, Tran SV, Weinstock JV, Gallini CA, Redding K, Margolskee RF, Osborne LC, et al. 2016. Tuft cells, taste-chemosensory cells, orchestrate parasite type 2 immunity in the gut. Science 351:1329–1333.

23. Huang J, Lee H-y, Zhao X, Han J, Su Y, Sun Q, Shao J, Ge J, Zhao Y, Bai X, et al. 2021. Interleukin-17D regulates group 3 innate lymphoid cell function through its receptor CD93. Immunity 54:673-686.e674.

24. Huson DH, Richter DC, Rausch C, Dezulian T, Franz M, Rupp R. 2007. Dendroscope: An interactive viewer for large phylogenetic trees. BMC Bioinformatics 8:460.

25. Kawaguchi M, Adachi M, Oda N, Kokubu F, Huang S-K. 2004. IL-17 cytokine family. Journal of Allergy and Clinical Immunology 114:1265–1273.

26. Kim E, Huh JR, Choi GB. 2024. Prenatal and postnatal neuroimmune interactions in neurodevelopmental disorders. Nature Immunology 25:598–606.

27. Kirilenko BM, Munegowda C, Osipova E, Jebb D, Sharma V, Blumer M, Morales AE, Ahmed A-W, Kontopoulos D-G, Hilgers L, et al. 2023. Integrating gene annotation with orthology inference at scale. Science 380:eabn3107.

28. Kolde R. 2019. Pheatmap: pretty heatmaps. R package version 1:726.

29. Kono T, Korenaga H, Sakai M. 2011. Genomics of fish IL-17 ligand and receptors: A review. Fish & Shellfish Immunology 31:635–643.

30. Kumar S, Stecher G, Suleski M, Hedges SB. 2017. TimeTree: A Resource for Timelines, Timetrees, and Divergence Times. Molecular Biology and Evolution 34:1812–1819.

31. Kwon H-K, Choi GB, Huh JR. 2022. Maternal inflammation and its ramifications on fetal neurodevelopment. Trends in Immunology 43:230–244.

32. Lammert CR, Frost EL, Bolte AC, Paysour MJ, Shaw ME, Bellinger CE, Weigel TK, Zunder ER, Lukens JR. 2018. Cutting Edge: Critical Roles for Microbiota-Mediated Regulation of the Immune System in a Prenatal Immune Activation Model of Autism. The Journal of Immunology 201:845–850.

33. Langley CA, Dietzen PA, Emerman M, Tenthorey JL, Malik HS. 2025. Antiviral Mx proteins have an ancient origin and widespread distribution among eukaryotes. Proceedings of the National Academy of Sciences 122:e2416811122.

34. Lee B, Kwon J-T, Jeong Y, Caris H, Oh D, Feng M, Davila Mejia I, Zhang X, Ishikawa T, Watson BR, et al. 2025. Inflammatory and anti-inflammatory cytokines bidirectionally modulate amygdala circuits regulating anxiety. Cell 188:2190-2202.e2115.

35. Lee Y, Ishikawa T, Lee H, Lee B, Ryu C, Davila Mejia I, Kim M, Lu G, Hong Y, Feng M, et al. 2025. Brain-wide mapping of immune receptors uncovers a neuromodulatory role of IL-17E and the receptor IL-17RB. Cell 188:2203-2217.e2217.

36. Leonardi I, Gao IH, Lin W-Y, Allen M, Li XV, Fiers WD, De Celie MB, Putzel GG, Yantiss RK, Johncilla M, et al. 2022. Mucosal fungi promote gut barrier function and social behavior via Type 17 immunity. Cell 185:831-846.e814.

37. Li Q, Li X, Lou Y, Xing Y, Yan W, Chen Y, Hu G, Song X, Liu Z, Yang T, Qian Y. 2026. The IL17REL gene encodes a decoy receptor of IL-17 family cytokines to control gut inflammation. Nature Immunology 27:225–235.

38. Little J, Chikina M, Clark NL. 2024. Evolutionary rate covariation is a reliable predictor of co-functional interactions but not necessarily physical interactions. Elife 12:RP93333.

39. Little J, Meyer GH, Grover A, Francette AM, Partha R, Arndt KM, Smith M, Clark N, Chikina M. 2025. ERC 2.0 - evolutionary rate covariation update improves inference of functional interactions across large phylogenies. bioRxiv:2025.2002.2024.639970.

40. Liu S, Song X, Chrunyk BA, Shanker S, Hoth LR, Marr ES, Griffor MC. 2013. Crystal structures of interleukin 17A and its complex with IL-17 receptor A. Nature Communications 4:1888.

41. Liu X, Sun S, Liu D. 2020. IL-17D: A Less Studied Cytokine of IL-17 Family. International Archives of Allergy and Immunology 181:618–623.

42. Madeira F, Madhusoodanan N, Lee J, Eusebi A, Niewielska A, Tivey ARN, Lopez R, Butcher S. 2024. The EMBL-EBI Job Dispatcher sequence analysis tools framework in 2024. Nucleic acids research 52:W521–W525.

43. Margolis SR, Wilson SC, Vance RE. 2017. Evolutionary Origins of cGAS-STING Signaling. Trends in Immunology 38:733–743.

44. McDonald NQ, Hendrickson WA. 1993. A structural superfamily of growth factors containing a cystine knot motif. Cell 73:421–424.

45. McGeachy MJ, Cua DJ, Gaffen SL. 2019. The IL-17 Family of Cytokines in Health and Disease. Immunity 50:892–906.

46. Murrell B, Moola S, Mabona A, Weighill T, Sheward D, Kosakovsky Pond SL, Scheffler K. 2013. FUBAR: A Fast, Unconstrained Bayesian AppRoximation for Inferring Selection. Molecular Biology and Evolution 30:1196–1205.

47. Murrell B, Weaver S, Smith MD, Wertheim JO, Murrell S, Aylward A, Eren K, Pollner T, Martin DP, Smith DM, et al. 2015. Gene-Wide Identification of Episodic Selection. Molecular Biology and Evolution 32:1365–1371.

48. Murrell B, Wertheim JO, Moola S, Weighill T, Scheffler K, Kosakovsky Pond SL. 2012. Detecting Individual Sites Subject to Episodic Diversifying Selection. PLOS Genetics 8:e1002764.

49. Musil M, Khan RT, Beier A, Stourac J, Konegger H, Damborsky J, Bednar D. 2020. FireProtASR: A Web Server for Fully Automated Ancestral Sequence Reconstruction. Briefings in Bioinformatics 22.

50. Nelson JC, Granato M. 2022. Zebrafish behavior as a gateway to nervous system assembly and plasticity. Development 149.

51. Nielsen R, Bustamante C, Clark AG, Glanowski S, Sackton TB, Hubisz MJ, Fledel-Alon A, Tanenbaum DM, Civello D, White TJ, et al. 2005. A Scan for Positively Selected Genes in the Genomes of Humans and Chimpanzees. PLOS Biology 3:e170.

52. OhAinle M, Kerns JA, Malik HS, Emerman M. 2006. Adaptive Evolution and Antiviral Activity of the Conserved Mammalian Cytidine Deaminase <i>APOBEC3H</i>. Journal of Virology 80:3853–3862.

53. Osborne LM, Brar A, Klein SL. 2019. The role of Th17 cells in the pathophysiology of pregnancy and perinatal mood and anxiety disorders. Brain, Behavior, and Immunity 76:7–16.

54. Pond SLK, Frost SDW, Muse SV. 2004. HyPhy: hypothesis testing using phylogenies. Bioinformatics 21:676–679.

55. Rambaut A, Drummond A. 2012. FigTree version 1.4. 0. In.

56. Reed MD, Yim YS, Wimmer RD, Kim H, Ryu C, Welch GM, Andina M, King HO, Waisman A, Halassa MM, et al. 2020. IL-17a promotes sociability in mouse models of neurodevelopmental disorders. Nature 577:249–253.

57. Reynolds JM, Angkasekwinai P, Dong C. 2010. IL-17 family member cytokines: Regulation and function in innate immunity. Cytokine & Growth Factor Reviews 21:413–423.

58. Saco A, Rey-Campos M, Rosani U, Novoa B, Figueras A. 2021. The Evolution and Diversity of Interleukin-17 Highlight an Expansion in Marine Invertebrates and Its Conserved Role in Mucosal Immunity. Frontiers in Immunology Volume 12 -2021.

59. Salvador AF, de Lima KA, Kipnis J. 2021. Neuromodulation by the immune system: a focus on cytokines. Nature Reviews Immunology 21:526–541.

60. Schneider C, O’Leary CE, Locksley RM. 2019. Regulation of immune responses by tuft cells. Nature Reviews Immunology 19:584–593.

61. Shin Yim Y, Park A, Berrios J, Lafourcade M, Pascual LM, Soares N, Yeon Kim J, Kim S, Kim H, Waisman A, et al. 2017. Reversing behavioural abnormalities in mice exposed to maternal inflammation. Nature 549:482–487.

62. Shultz AJ, Sackton TB. 2019. Immune genes are hotspots of shared positive selection across birds and mammals. Elife 8:e41815.

63. Sievers F, Wilm A, Dineen D, Gibson TJ, Karplus K, Li W, Lopez R, McWilliam H, Remmert M, Söding J, et al. 2011. Fast, scalable generation of high-quality protein multiple sequence alignments using Clustal Omega. Molecular Systems Biology 7:MSB201175.

64. Sutton CE, Mielke LA, Mills KHG. 2012. IL-17-producing γδ T cells and innate lymphoid cells. European Journal of Immunology 42:2221–2231.

65. von Moltke J, Ji M, Liang H-E, Locksley RM. 2016. Tuft-cell-derived IL-25 regulates an intestinal ILC2–epithelial response circuit. Nature 529:221–225.

66. Wilson SC, Caveney NA, Yen M, Pollmann C, Xiang X, Jude KM, Hafer M, Tsutsumi N, Piehler J, Garcia KC. 2022. Organizing structural principles of the IL-17 ligand–receptor axis. Nature 609:622–629.

67. Wu B, Jin M, Zhang Y, Wei T, Bai Z. 2011. Evolution of the IL17 receptor family in chordates: a new subfamily IL17REL. Immunogenetics 63:835–845.

68. Yang L, Huh JR, Choi GB. 2023. One messenger shared by two systems: How cytokines directly modulate neurons. Current Opinion in Neurobiology 80:102708.

69. Yang Z. 2007. PAML 4: Phylogenetic Analysis by Maximum Likelihood. Molecular Biology and Evolution 24:1586–1591.

70. Yuan L, Huang J, Chen J, Xie T, Wang G, Xie B, Qin L, Chen Y, Zhong X, Zhao Z, et al. 2025. Ciliated cell–derived IL-17D restrains allergic asthma through controlling monocyte recruitment. Journal of Experimental Medicine 222.

71. Zhang X, Angkasekwinai P, Dong C, Tang H. 2011. Structure and function of interleukin-17 family cytokines. Protein & Cell 2:26–40.

