## Supplementary material for "Genetic comparisons of interleukin-17 reveal a framework for complex signaling evolution": IL17a busted analysis.txt

Analysis Description
--------------------
BUSTED (branch-site unrestricted statistical test of episodic
diversification) uses a random effects branch-site model fitted jointly
to all or a subset of tree branches in order to test for alignment-wide
evidence of episodic diversifying selection. Assuming there is evidence
of positive selection (i.e. there is an omega > 1), BUSTED will also
perform a quick evidence-ratio style analysis to explore which
individual sites may have been subject to selection. v2.0 adds support
for synonymous rate variation, and relaxes the test statistic to 0.5
(chi^2_0 + chi^2_2). Version 2.1 adds a grid search for the initial
starting point. Version 2.2 changes the grid search to LHC, and adds an
initial search phase to use adaptive Nedler-Mead. Version 3.0 implements
the option for branch-site variation in synonymous substitution rates.
Version 3.1 adds HMM auto-correlation option for SRV, and binds SRV
distributions for multiple branch sets. Version 4.0 adds support for
multiple hits (MH), ancestral state reconstruction saved to JSON, and
profiling of branch-site level support for selection / multiple hits.
Version 4.2 adds calculation of MH-attributable fractions of
substitutions. Version 4.5 adds an 'error absorption' component
[experimental] 

- __Requirements__: in-frame codon alignment and a phylogenetic tree (optionally annotated
with {})

- __Citation__: *Gene-wide identification of episodic selection*, Mol Biol Evol.
32(5):1365-71, *Synonymous Site-to-Site Substitution Rate Variation
Dramatically Inflates False Positive Rates of Selection Analyses: Ignore
at Your Own Peril*, Mol Biol Evol. 37(8):2430-2439

- __Written by__: Sergei L Kosakovsky Pond

- __Contact Information__:

- __Analysis Version__: 4.5


>code => Universal
/home/datamonkey/datamonkey-js-server/production/app/busted/output/67210bf53669ec76d3b01916.tre
/home/datamonkey/datamonkey-js-server/production/app/busted/output/67210bf53669ec76d3b01916.tre

>Loaded a multiple sequence alignment with **29** sequences, **150** codons, and **1** partitions from `/home/datamonkey/datamonkey-js-server/production/app/busted/output/67210bf53669ec76d3b01916`

>branches => All

>srv => Yes
The number omega rate classes to include in the model (permissible range = [1,10], default value = 3, integer): 
>rates => 3

>multiple-hits => None
The number alpha rate classes to include in the model [1-10, default 3] (permissible range = [1,10], default value = 3, integer): 
>syn-rates => 3

>error-sink => No
The number of points in the initial distributional guess for likelihood fitting (permissible range = [1,10000], default value = 250, integer): 
>grid-size => 250
The number of initial random guesses to 'seed' rate values optimization (permissible range = [1,25], default value = 1, integer): 
>starting-points => 1


### Branches to test for selection in the BUSTED analysis
* Selected 55 branches to test in the BUSTED analysis: `AOTUS_NANCYMAAE, CEBUS_IMITATOR, SAPAJUS_APELLA, Node10, SAIMIRI_BOLIVIENSIS, Node9, Node7, CALLITHRIX_JACCHUS, Node6, CARLITO_SYRICHTA, LEMUR_CATTA, PROPITHECUS_COQUERELI, Node19, MICROCEBUS_MURINUS, Node18, NYCTICEBUS_COUCANG, OTOLEMUR_GARNETTII, Node23, Node17, Node15, Node5, GORILLA_GORILLA_GORILLA, HOMO_SAPIENS, Node28, PONGO_ABELII, Node27, HYLOBATES_MOLOCH, NOMASCUS_LEUCOGENYS, SYMPHALANGUS_SYNDACTYLUS, Node34, Node32, Node26, Node4, COLOBUS_ANGOLENSIS_PALLIATUS, PILIOCOLOBUS_TEPHROSCELES, Node38, RHINOPITHECUS_BIETI, RHINOPITHECUS_ROXELLANA, Node42, TRACHYPITHECUS_FRANCOISI, Node41, Node37, Node3, CHLOROCEBUS_SABAEUS, Node2, PAPIO_ANUBIS, THEROPITHECUS_GELADA, Node47, Node1, CERCOCEBUS_ATYS, MACACA_FASCICULARIS, MACACA_NEMESTRINA, Node52, Node50, MACACA_MULATTA`


### Obtaining branch lengths and nucleotide substitution biases under the nucleotide GTR model

>kill-zero-lengths => No
* Log(L) = -2104.86, AIC-c =  4336.35 (63 estimated parameters)
* 1 partition. Total tree length by partition (subs/site)  0.651

### Obtaining the global omega estimate based on relative GTR branch lengths and nucleotide substitution biases
* Log(L) = -2059.80, AIC-c =  4261.92 (70 estimated parameters)
* 1 partition. Total tree length by partition (subs/site)  0.655
* non-synonymous/synonymous rate ratio for *test* =   0.4337

### Improving branch lengths, nucleotide substitution biases, and global dN/dS ratios under a full codon model
* Log(L) = -2058.40, AIC-c =  4259.12 (70 estimated parameters)
* non-synonymous/synonymous rate ratio for *test* =   0.4148

### Performing the full (dN/dS > 1 allowed) branch-site model fit
